# Recurrent activity within microcircuits of macaque dorsolateral prefrontal cortex tracks cognitive flexibility

**DOI:** 10.1101/2023.09.23.559125

**Authors:** Suzanne O Nolan, Patrick R Melugin, Kirsty R Erickson, Wilson R Adams, Zahra Z Farahbakhsh, Colleen E Mcgonigle, Michelle H Kwon, Vincent D Costa, Christopher C Lapish, Troy A Hackett, Verginia C Cuzon Carlson, Christos Constantinidis, Kathleen A Grant, Cody A Siciliano

**Author notes:** **Corresponding authors:** Cody A Siciliano ✉ Kathleen A Grant ✉.

## Abstract

Human and non-human primate data clearly implicate the dorsolateral prefrontal cortex (dlPFC) as critical for advanced cognitive functions^1,2^. It is thought that intracortical synaptic architectures within dlPFC are the integral neurobiological substrate that gives rise to these processes, including working memory, inferential reasoning, and decision-making^3–7^. In the prevailing model, each cortical column makes up one fundamental processing unit composed of dense intrinsic connectivity, conceptualized as the ‘canonical’ cortical microcircuit^3,8^. Each cortical microcircuit receives sensory and cognitive information from a variety of sources which are represented by sustained activity within the microcircuit, referred to as persistent or recurrent activity^4,9^. Via recurrent connections within the microcircuit, activity can propagate for a variable length of time, thereby allowing temporary storage and computations to occur locally before ultimately passing a transformed representation to a downstream output^4,5,10^. Competing theories regarding how microcircuit activity is coordinated have proven difficult to reconcile *in vivo* where intercortical and intracortical computations cannot be fully dissociated^5,9,11,12^. Here, we interrogated the intrinsic features of isolated microcircuit networks using high-density calcium imaging of macaque dlPFC *ex vivo*. We found that spontaneous activity is intrinsically maintained by microcircuit architecture, persisting at a high rate in the absence of extrinsic connections. Further, using perisulcal stimulation to evoke persistent activity in deep layers, we found that activity propagates through stochastically assembled intracortical networks, creating predictable population-level events from largely non-overlapping ensembles. Microcircuit excitability covaried with individual cognitive performance, thus anchoring heuristic models of abstract cortical functions within quantifiable constraints imposed by the underlying synaptic architecture.

## Results & Discussion

Contemporary knowledge of cortical microcircuits relies heavily on studies in non-primate species, where *ex vivo* brain slice preparations allow the cytoarchitectural organization and synaptic weights dictating intracortical dynamics to be interrogated in the absence of intercortical and sensory inputs^13–15^. While these studies have been critical to our understanding of the cortical microcircuitry of sensory cortices, the dlPFC is an evolutionary specialization distinct to primates without a direct homolog in other mammals^16,17^. As such, direct investigation of the intrinsic properties of intracortical dlPFC microcircuits in primates represents a missing link in the chain of evidence connecting synaptic features at the single-cell level with the cognitive processes that the dlPFC subserves. This work establishes a paradigm for assessing intrinsic features of intracortical microcircuits in acute primate brain slices to explore three overarching questions: 1) to what degree does the intrinsic architecture of the dlPFC propagate activity in the absence of ongoing extrinsic inputs, 2) for a given discrete input, does network activity arise from dedicated, stable groups of cells or from a distributed and flexible network, and 3) do the static features of intracortical excitability observed *ex vivo* covary with individual differences in cognitive performance.

To interrogate microcircuit interactions that govern network-level properties, it is necessary to observe the activity of many cells simultaneously, ideally without sampling bias towards genetically- or morphologically-defined subpopulations. To achieve high-density, spatiotemporally resolved recordings of randomly sampled cells to permit unbiased interrogation of population dynamics, acute *ex vivo* brain slices from adult rhesus macaques containing dlPFC area 46 were loaded with the cell-permeable fluorescent calcium dye fluo-4 acetoxymethyl ester (fluo-4 AM) **(Figure S1A)**. Using necropsy and MRI-guided tissue preparation procedures optimized for obtaining viable slices with high anatomical repeatability^18,19^, slices displayed high loading efficiency with cells clearly visible with as little as a one-minute incubation in 5 µM of fluo-4 AM **(Figure S1B)**. Importantly, fluo-4 AM is cell-permeable and non-fluorescent until cleavage of the AM tag by intracellular esterases, which renders the dye cell-impermeable and activates its calcium-dependent fluorescence, thus trapping the active form within the cell. Because this process requires active esterases, loading efficiency also provides a within-sample, online verification of cell viability^20,21^. A 30-minute loading period was sufficient to visualize cellular processes and produced dense loading throughout the slice **(Figure S1B-E)**. Co-staining with DAPI confirmed that cellular features aligned with nuclei as expected **(Figure S1F)**. Using high-speed laser scanning confocal microscopy, we visualized temporally-resolved calcium dynamics in hundreds of cells simultaneously over a 762 µm^2^ field of view.

To systematically evaluate the degree to which a temporally and spatially discrete stimulation produces persistent activity, a bipolar stimulating electrode was placed superficially along the bank of the principal sulcus while calcium recordings were obtained distally in the deep pyramidal layers (**Figure 1A**-**D**). This configuration allows for input-output relationships to be directly assessed in the absence of extrinsic connections, which necessitates that any polysynaptic transient changes in activity must be propagated through feedback and/or feedforward connections within the slice (referred to throughout as recurrent). By recording distal to the stimulating electrode, recurrent activity can be assessed downstream of the direct depolarizations induced by the electrical field potential generated between the poles of the electrode. Though stimulation cannot be equated with endogenous inputs occurring in an intact system, any propagation of activity downstream of the initial event must conform to the synaptic architecture and intrinsic constraints imposed by the intracortical microcircuitry. Using this approach, we first examined how recurrent activity patterns arise from a single pulse stimulation across a range of intensities (n=8 slices from 8 subjects, 2-10 mA single pulse stimulations). We found that single pulse stimulation was sufficient to produce robust transients in the bulk fluorescence signal (mean intensity over time across the field of view) which scaled with increasing input intensities (**Figure 1E**). Intensity-dependent increases were observed in both peak amplitude as well as the area under the curve of the decay phase, measured in a sliding 2s window beginning one frame after the peak fluorescence intensity for each trace (**Figure 1G**-**H**). Importantly, the stimulation duration (4 ms) and the temporal kinetics of the sensor (rise time = approximately 40 ms^22^) are far shorter than the duration of the observed signals, indicating propagation of activity via multi-synaptic events and/or persistent changes in basal activity patterns. Moreover, the loading efficiency, as assessed by the number of cells segmented in each field of view, did not covary with the amplitude of the whole field response (**Figure S1G**), confirming that these signals do not simply reflect loading efficiency. The input-output curves for both signal amplitude and area under the curve of the decay phase displayed a sigmoidal shape over scaling intensities. Indeed, over a relatively restricted range of input intensity (5-fold), both curves scaled to asymptote and displayed similar half-maximal intensities of roughly 5 mA (**Figure 1G**-**H**). These results indicate that propagation of intracortical activity can be readily evoked in this preparation and that, despite the exogenous nature of the input stimulation, disparate input parameters produce distinct and predictable population-level responses.

**Figure 1.**
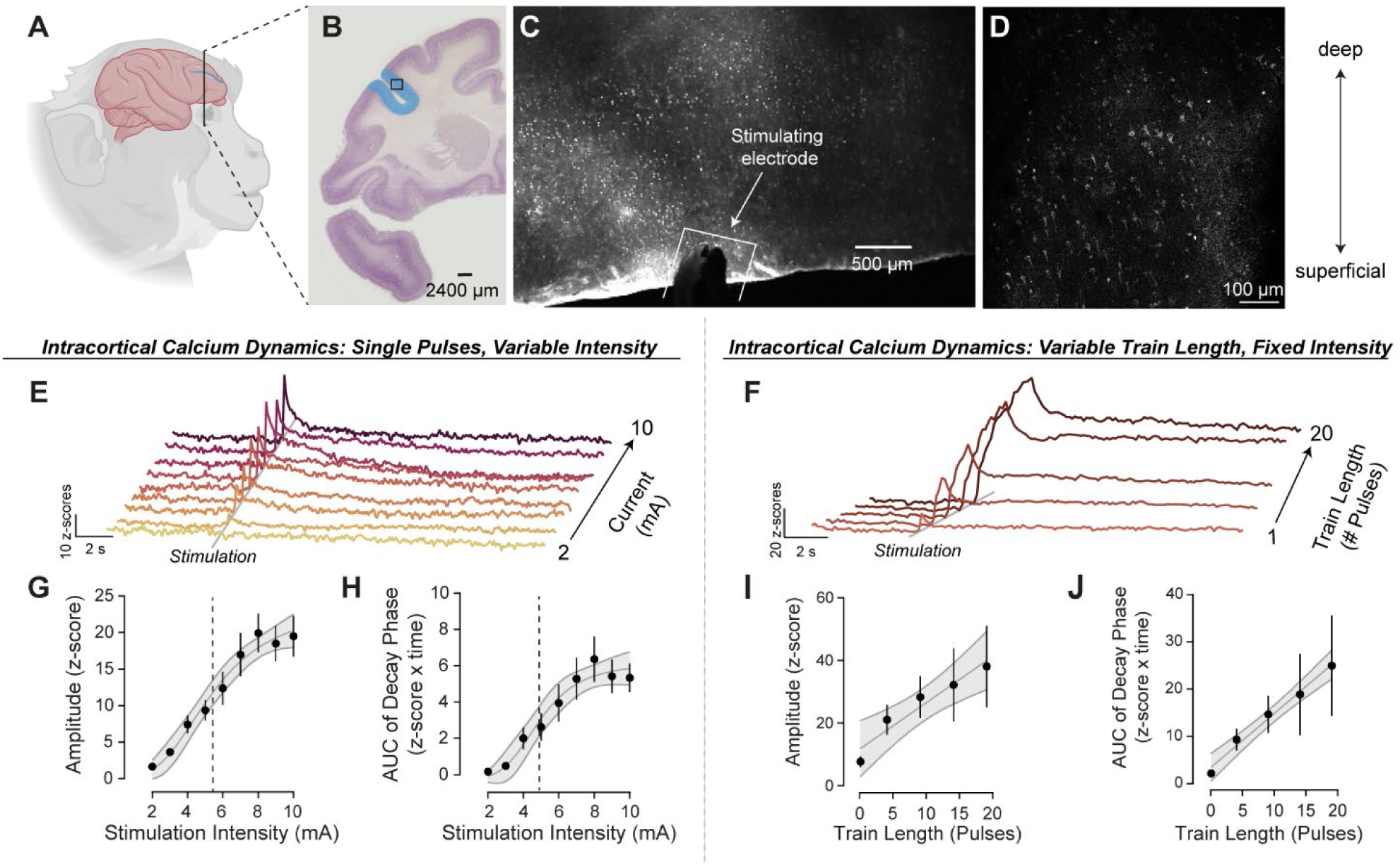
Localized peri-sulcal stimulation elicits far-field representations in population-level signals. **(A)** *Ex vivo* brain slices from 8 adult rhesus macaques were prepared immediately post-mortem from tissue blocks sectioned in the coronal plane. **(B)** All recordings were performed in Brodmann’s area 46 of the dorsolateral prefrontal cortex, indicated in blue (atlas image was reproduced from BrainMaps.Org). **(C)** To deliver a localized input stimulus to the superficial cortex without inducing an electrical potential across layers, a twisted bipolar electrode was placed peri-sulcal with contacts parallel to the bank of the principal sulcus. The image depicts stimulation placement and gross anatomical features, which were assessed using a low magnification air objective (a water dipping objective was used for all experimental recordings). **(D)** Cells were bulk loaded with the cell-permeable fluorescent calcium indicator fluo-4 AM and *ex vivo* calcium dynamics were observed across a 762 µm^2^ field of view at 15.14 fps. The image shows a standard deviation T-projection from a representative field of view. **(E)** Single pulses with varying amplitude were delivered to test the input-output relationship across input intensities while all other parameters were held constant (monophasic, 4 ms pulse width). Representative brightness-over-time traces of bulk fluorescence demonstrate far-field responses evoked by peri-sulcal single pulse stimulations (2-10 mA constant current, ascending order, stimulation onset at 5 seconds). **(F)** In a parallel experiment, input-output relationships were examined for stimulations of varying train lengths while all other parameters were held constant (1-20 pulses, ascending order, 20 Hz). **(G)** The amplitude of evoked bulk fluorescence activity scales nonlinearly with stimulation intensity and was better described by a four-parameter symmetrical logistic function than by a linear fit (extra sum-of-squares F test, F_(1,6)_ = 9.28, p< 0.05 for fit comparison; R^2^ = 0.98 for nonlinear fit). Vertical line denotes the EA_50_ value (5.42 mA, [95% CI: 4.51, 10.11]). **(H)** Area under the curve (AUC) of the decay phase, measured in a sliding window starting one frame after the peak of each trace, also scales nonlinearly as a function of input intensity (extra sum-of-squares F test, F_(1,6)_ = 12.05, p< 0.05 vs linear fit, R^2^ = 0.96 for nonlinear fit). Vertical line denotes the EA_50_ value (4.91 mA [95% CI: 4.14, 7.13]). **(I)** Across stimulations of increasing train length, there was a linear increase in the amplitude of the bulk fluorescence response (extra sum-of-squares F test, F_(1,2)_ = 17.98, p > 0.05 vs linear fit, R^2^ = 0.92 for linear fit). **(J)** Similarly, the AUC of the decay phase scaled linearly with train length (extra sum-of-squares F test, F_(1,2)_ = 1.92, p > 0.05 vs linear, R^2^ = 0.99 for linear fit). Error bars indicate SEM. For curve fit data, error bands indicate 95% confidence intervals of best fit values.

In a parallel experiment (n=5 slices from 5 subjects), using the same recording configuration, we first evoked activity using a single pulse stimulation matched to the half-maximal parameters observed in the stimulation intensity curve (5 mA, all parameters identical). We then examined the input-output relationship across an orthogonal parameter axis by delivering stimulations of fixed intensity with increasing train lengths (1 – 20 pulses, 20 Hz, 5 mA throughout) (**Figure 1F**). The amplitudes and area of the decay phase evoked by the single pulse, 5 mA stimulation did not differ between experiments (unpaired t-test_(1,11)_=0.66, p = 0.52 for peak amplitude; unpaired t-test_(1,11)_=0.86, p = 0.41 for AUC), confirming that this recording strategy allows repeatable input-output relationships to be observed and compared across slices. The input-output scaling for repeated pulses differed markedly from the intensity curve. Both amplitude and AUC of the decay phase increased linearly with increasing train lengths, and scaled to magnitudes well beyond the plateau of the intensity curve without displaying any apparent non-linearity (**Figure 1I**-**J**). These results indicate inputs received in close temporal succession, 50 ms apart in this case, produce summation in downstream activity. This temporal summation is cumulative, scaling linearly even when inputs arrive over a relatively extended total timeframe (1s between the first and last pulse in the 20 pulse condition).

We next extracted activity traces corresponding to single cells from the recordings. Notably, on the single-cell level, there was a striking degree of spontaneous activity in the absence of exogenous stimulation (Video S1) **(Figure S2A-B)**. This is in stark contrast to previous reports that rodent PFC displays high rates of spontaneous activity *in vivo* but is largely silent when observed in the absence of extrinsic inputs^13,23,24^, and demonstrates that dlPFC is self-sufficient in maintaining active networks. To determine how inputs are integrated across individual cells to give rise to the population level events described above, we next visualized activity aligned to stimulation onset for the 2,189 cells recorded across the intensity curve (**Figure 2A**). Single-cell activity traces were then thresholded to classify the cells according to responsivity (excited, inhibited, no response). Separation into excited or inhibited subpopulations illustrated stimulation-evoked divergent activity between the subpopulations and unclassified (no response) cells as expected, and demonstrated that this divergent activity persisted well beyond the window used for classification (**Figure 2B**). Comparing the distribution of excited, inhibited, and non-responsive cells across stimulations showed moderate increases in the number of excited and inhibited cells as the stimulation intensity increased (**Figure 2C**). Indeed, increasing stimulation intensity led to a linear rise in the proportion of the population recruited (**Figure 2D**). Further, analysis of individual amplitudes for each cell across the intensity curve showed that the cells classified as having no response showed minimal-to-no input-dependent scaling, corroborating that the thresholds set for classification effectively captured separate subpopulations **(Figure S2C-J).** To further validate the integrity of these measures, we also confirmed that both excitatory and inhibitory responses were blocked by the calcium chelator BAPTA-AM **(Figure S2K-L)**. Examination of single-cell responses over the input-output curve for increasing train length (n = 549 cells) again demonstrated that persistent excitatory and inhibitory changes in the activity of single cells were evident well beyond the initial stimulation-evoked response (**Figure 2E**-**F**). The proportion of the population recruited by the stimulation scaled with increasing train lengths (**Figure 2G**). In contrast to the intensity curve, where proportions of the population recruited scaled linearly over a limited range, increasing the train length led to robust and distinctly nonlinear increases in the proportion of cells responsive to the stimulation (**Figure 2H**). Thus, though there was clear scaling in the amplitude of the bulk responses over both curves (**Figure 1E**-**F**), the input-output relationship was disparate. Together these results suggest that temporal summation of inputs is capable of recruiting a larger downstream network than is possible with any single input, regardless of intensity. We next sought to determine if the input-output relationships described above are mediated by specific networks of cells. The networks within a cortical microcircuit and their contribution to neural processes are the subject of long-standing debate. Though the terminology has taken many forms over the decades, a repeated theme is whether networks of co-active cells, which we will refer to here as ensembles, are composed of dedicated versus labile members. Theories have spanned the spectrum of this dichotomy^25–28^. On one extreme, the base unit of cognitive information is stored in a physical location, or engram, within an ensemble of cells that each have a dedicated contribution to the ensemble dynamics when activated^25^. On the other extreme, ensembles are functionalized through a population-level code where any given output can be achieved by distinct combinations of cellular members such that contributions of each are interchangeable, and no single component is indispensable^27,28^. *In vivo* recordings have demonstrated the existence and importance of ensembles in sensory and cognitive representations^29,30^; however, at any given time, an unknown number of ensembles are active, due to encoding of sensory stimuli and internal cognitive processes which are constantly co-occurring *in vivo*. This complexity, along with a multitude of technical limitations in recording from identified cells, has prevented progress towards reconciling competing theories of ensemble membership stability in the dlPFC.

**Figure 2.**
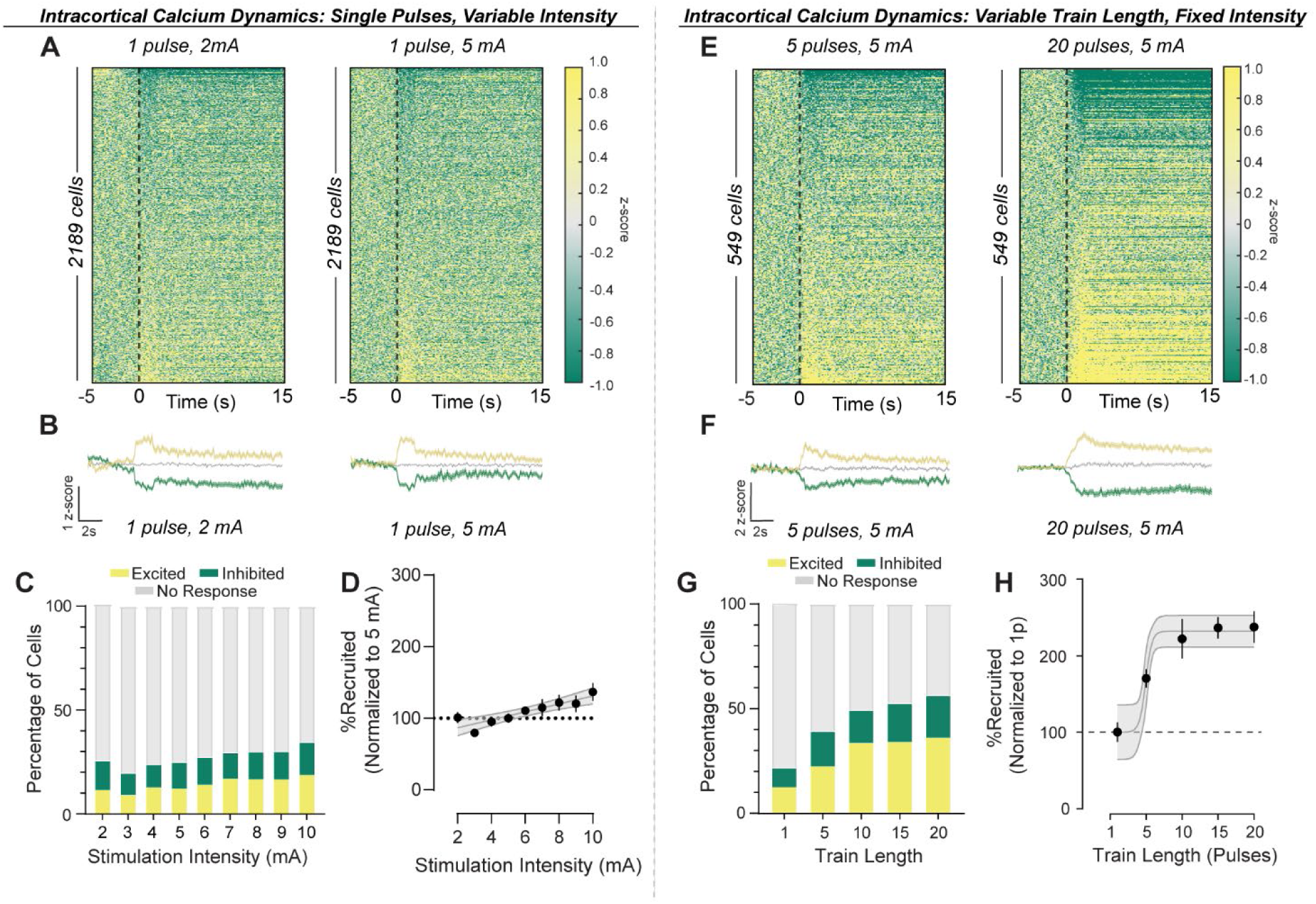
Evoked intracortical dynamics are enabled via recursion within scalable ensembles. **(A)** Heatmaps of single-cell dynamics for single pulse stimulations (2 and 5 mA) with stimulation onset indicated by the dotted black line. **(B)** Activity over time traces for population averages among cells classified as excited, inhibited, or not responsive to 2 mA and 5 mA single pulse stimulations. **(C)** Distributions of response types (Excited, Inhibited, No response) as a percent of the population differs across stimulation intensities (chi-squared test, χ^2^ (16) =197.8, p < 0.0001). **(D)** The percentage of cells recruited by the stimulation scales linearly with increasing stimulation intensity (extra sum-of-squares F test, F_(2,68)_ = 0.62, p > 0.05 vs linear fit, R^2^ = 0.24 for linear fit; F_(1,70)_ = 22.54, p < 0.0001 vs zero). **(E)** Representative heatmaps of single-cell dynamics evoked by multiple pulses (5 pulses and 20 pulses at 5 mA). **(F)** Activity over time traces for population averages among cells classified as excited, inhibited, or no response to 5 pulse and 20 pulse stimulations. **(G)** Proportion of the population recruited increases across stimulations with increasing train length (chi-squared test, χ^2^ (8) =179.0, p < 0.0001). **(H)** Increasing train length leads to nonlinear increases in ensemble size (extra sum-of-squares F test, F_(1,22)_ = 8.22, p < 0.05 vs linear fit, R^2^ = 0.68 for nonlinear fit). Error bars indicate SEM. For curve fit data, error bands indicate 95% confidence intervals of best fit values.

To explore the range of possibilities permitted by the intrinsic connectivity of the dlPFC, we analyzed the ensembles evoked across the input-output curves of stimulation intensity and train length with respect to cellular identities. Activity traces, detailed in Figure 1, were extracted from a single concatenated, co-registered video of all stimulation replicates per slice/subject. Thus, any possible methodological weights on detection thresholds were applied equally across all stimulations and each cell in the dataset received an unambiguous, binary classification indicating the presence or absence of activity at each stimulation replicate.

Throughout both experiments, the stimulating electrode remained in a static location, allowing for observations of ensembles evoked by a stimulus (***S***) with known features in the absence of any non-recurrent inputs to the network. Stimulations were delivered at least 3 minutes apart, thus ensembles evoked by each iteration (***x***) of **S** were considered as the representation of ***S^x^* (Figure 3A)**. The population of cells recorded throughout the experiment are referred to as ***S^Δ^***, and probabilities were expressed as a proportion of the number of cells in the ***S^Δ^*** population. We first determined the shared versus unique membership between the ensembles evoked by the first, lowest intensity stimulation (***S^A^***, corresponding to the 2 mA stimulation shown in Figure 1) and the stimulation which produced a half-maximal population-level response (***S^A^***→***S^D^***), and then again between the half-maximal stimulation and the final, highest intensity stimulation which was in the asymptotic portion of the input-output curve (***S^D^*** → ***S^I^***). A Sankey diagram depicting the percentage of ***S^Δ^*** recruited to each ensemble and the proportion of crossover between each, revealed that although ensemble size was essentially identical for ***S^A^ → S^D^***(26% and 25% of ***S^Δ^***, respectively), greater than 50% of the ***S^D^*** ensemble represented unique members which were not recruited in ***S^A^* (Figure 3B)**. At the highest intensity stimulation, the ensemble size scaled to 35%, and again more than 50% of the membership was unique from ***S^D^***→***S^I^***. Visualizing membership crossover between one, ten, and twenty pulse stimulations again showed a striking degree of membership instability (**Figure 3C**). Performing the same analysis within only the cells which were members of the ***S^A^*** ensemble yielded values strikingly similar to the full population with 25% and 36% of the ***S^A^*** ensemble being recruited to ***S^D^*** and ***S^I^***, respectively (**Figure 3D**). Dividing cells by whether they displayed excitatory or inhibitory responses revealed that a similar degree of turnover occurred within each response class (**Figure 3D-E**). These results indicate that in the absence of ongoing inputs, regardless of whether a broad or directionality-specific membership criterion is applied, recursive activity originating from a fixed spatial location within the dlPFC microcircuit propagates through predominantly labile ensembles.

**Figure 3.**
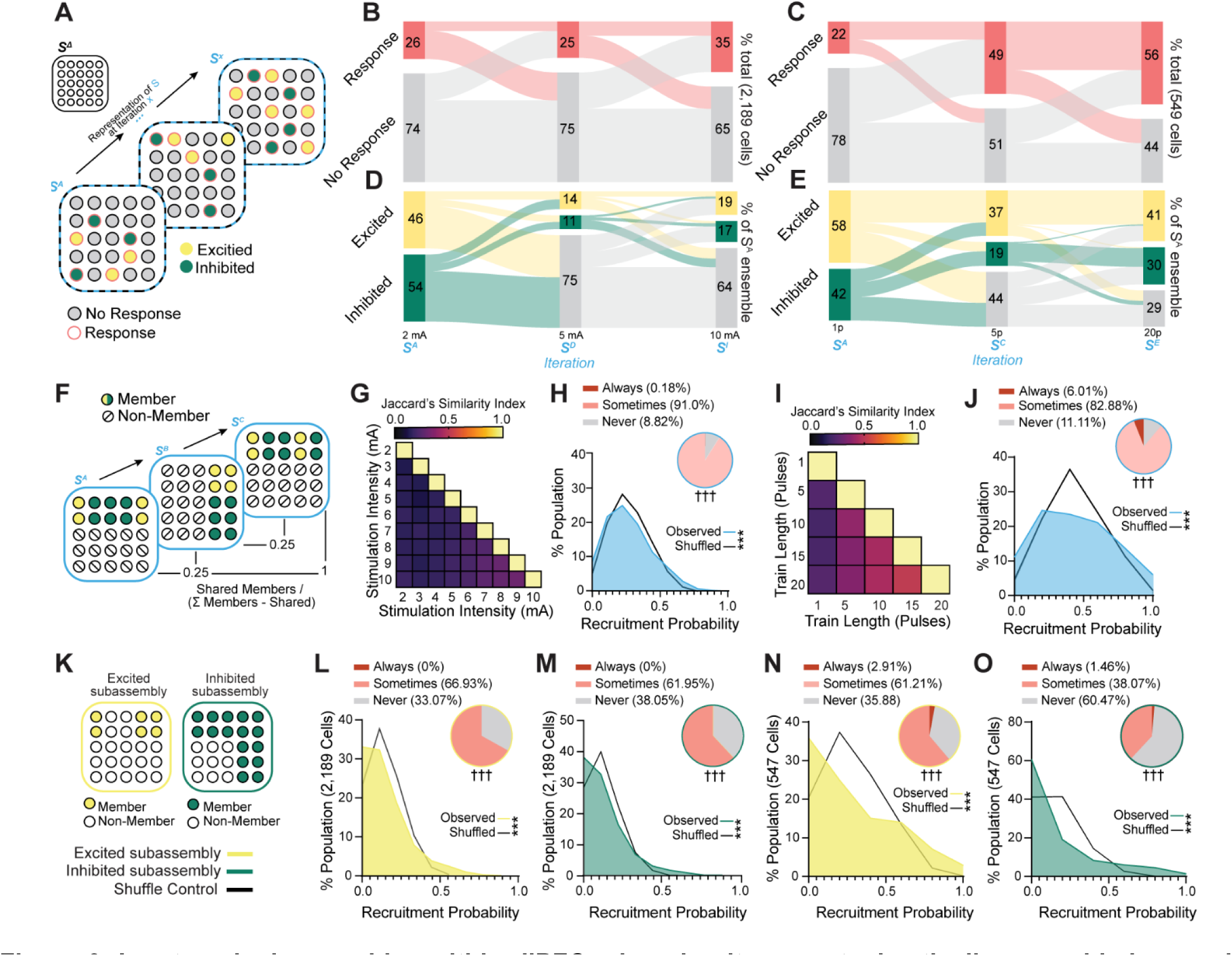
Input-evoked ensembles within dlPFC microcircuitry are stochastically assembled around a minority of core members. **(A)** Diagram of possible ensemble dynamics underlying the population level signals reported in Figure 1. *S^Δ^* refers to the population observed throughout the recordings regardless of activity, and *S^x^* refers to the ensemble activated by a stimulus *S* at a given iteration *x*. In this example, there is a gradual increase in the proportions of the population participating in the ensemble over iterations, but each iteration is represented by mostly unique members. **(B)** Sankey diagram showing retention/crossover of cells categorized as responsive vs not response to stimulation for the lowest intensity/first stimulation to the half-maximal/fifth stimulation (*S^A^* → *S^D^*) and from the half-maximal intensity stimulation to the final stimulation (*S^D^* → *S^I^*). **(C)** Sankey diagram showing retention/crossover of cells categorized as responsive vs no response to stimulation for the single pulse stimulation ten pulse/third stimulation in the train length curve dataset and from the ten-pulse stimulation to the twenty-pulse stimulation. **(D)** Sankey diagram of the same ensemble iterations as depicted in panel B, from the stimulation intensity curve experiment, but only including cells which were recruited to *S^A^* and with response directionality denoted. **(E)** Sankey diagram of the same ensemble iterations as depicted in panel C, from the train length curve experiment, but only including cells which were recruited by the first stimulation in the experiment and with response directionality denoted. **(F)** Diagram depicting potential ensemble dynamics that could underlie the high degree of turnover observed in both datasets (panels B-E). In the example, one set of cells switches in and out of the ensemble, resulting in a low similarity index between some of the iterations while others are identical (Jaccard’s index of 1). **(G)** Heatmaps of Jaccard’s similarity index for the 36 comparisons resulting from pairwise analysis of the ensemble memberships across the 9 iterations/stimulations intensities for the intensity curve dataset. An index of 1 indicates that members are identical, while an index of 0 indicates that members are non-overlapping. **(H)** Distribution of recruitment probability over the 9 iterations of ensemble *S* (number of times recruited to the ensemble / number of iterations). Shuffle distribution shown is the first of 10,000 reshuffles, for illustrative purposes (see Figure S6B for reshuffle statistics and distributions). Inset: cells were classified based on whether they were recruited to all of the ensemble iterations (‘Always’), at least one but not all (‘Sometimes’), or were not recruited to any of the ensemble iterations (‘Never’) (see Figure S6C for reshuffle comparisons). **(I)** Heatmaps of Jaccard’s similarity index for the 10 comparisons resulting from pairwise analysis of the ensemble memberships across the 5 iterations/stimulations from the variable train length dataset. **(J)** Distribution of recruitment probability over the 5 iterations of the ensemble evoked by stimulations of increasing train length (number of times recruited to the ensemble / number of iterations). The shuffle distribution shown is the first of 10,000 reshuffles, for illustrative purposes (see Figure S6D for reshuffle statistics and distributions). Inset: cells were classified as ‘Always’, ‘Sometimes’, or ‘Never’ based on whether they were recruited to all of the ensemble iterations, at least one but not all, or were not recruited to any of the ensemble iterations (see Figure S6D for reshuffle comparisons). **(K-O)** To determine if weighting of recruitment probability across the population was related to whether ensemble members were recruited through excitation or inhibition, datasets were reanalyzed twice using modified definitions of ensemble membership; once in which only stimulation-evoked increases in activity were considered for membership (excited subassembly) and once in which only stimulation-evoked decreases in activity were considered for membership (inhibited subassembly). **(K)** Diagram illustrating how differential weighting of recruitment probability across excited and inhibited cells can affect subassembly membership. The memberships depicted are from the hypothetical scenario depicted in panel F. **(L)** Recruitment probability over the 9 iterations in the stimulation intensity experiment when only excitatory responses are considered. Inset: associated classifications into Always excited, Sometimes exited, and Never excited. **(M)** Recruitment probability for the stimulation intensity experiment when only inhibitory responses are considered. Inset: associated classifications into Always inhibited, Sometimes inhibited, and Never inhibited. **(N)** Recruitment probability for the train length experiment when only the excitatory subassembly of the ensemble is considered. Inset: associated classifications. **(O)** Recruitment probability for the train length experiment when only the inhibitory subassembly of the ensemble is considered. Inset: associated classifications. *** median Kolmogorov-Smirnov D for observed vs shuffled from 10,000 reshuffles > critical value when α=0.001; ^†††^ median *X*^2^ value for observed vs shuffled from 10,000 reshuffles > critical value when α=0.001.

We next considered whether these dynamics were reflective of 1) a flexible but predictable ensemble, with the same sets of cells switching in and out at each iteration of ***S*,** or 2) probabilistic recruitment of unique members at each iteration such that a high degree of turnover would be observed at any intersection of ***S*** ensembles. Therefore, we determined the set similarity expressed as Jaccard’s similarity index, where values of 1 and 0 represent identical vs non-overlapping memberships for a given comparison, between each of the ensemble iterations (**Figure 3F**). The resulting set similarity maps clearly support the second scenario: across the 46 total comparisons resulting from pairwise analyses within each dataset similarity coefficients ranged from 0.12 to 0.47, demonstrating a high degree of turnover between any intersection of ***S*** ensembles **(Figure 3G, I)**.

These findings strongly suggest that the synaptic architecture of dlPFC microcircuitry is organized such that similar population-level signals can arise from probabilistically recruited networks.

Given the surprising degree of apparent instability, we next sought to determine whether ensemble recruitment was entirely probabilistic, with all cells having an equal likelihood of recruitment to the ensemble or, alternatively, if recruitment was weighted towards or against specific cells. To test these possibilities, a matrix was formed with columns equal to the number of cells in the dataset and rows equal to the number of iterations/stimulations. Each value in the matrix was a 0 or 1 indicating whether the cell was a member of the ensemble at each iteration. To determine the distribution of memberships that would be expected if recruitment probability was equally weighted across all cells, a shuffled dataset was created by randomizing the location of values in the matrix. Thus, the shuffled dataset had the same number of cells and stimulations, as well as the same total number of ensemble memberships, but any influence of cell identity on probability of ensemble membership across iterations was due to chance. Recruitment probability per cell [number of times included in the ensemble/number of times tested] was calculated for the observed and shuffled datasets, and the distributions of recruitment probability were compared across 10,000 rounds of reshuffling **(Figure S3A)**. We found that the distribution of recruitment probability to the ***S*** ensembles was consistently different than the shuffled control distributions, indicating that recruitment probability was unequally weighted across the population (**Figure 3H**, **Figure S3B)**.

To provide a more intuitive readout of how recruitment probability was weighted across the population, we categorized each cell based on whether it was activated at each iteration of ***S*** into three groups: those that were members at all iterations of ***S*** (termed “Always”), those that were not members of any of the ***S*** ensembles (termed “Never”), and those that were members and non-members of at least one ***S*** ensemble (termed “Sometimes”) (**Figure 3H**). The mean recruitment probability across iterations of ***S*** was 0.2731, thus the probability 𝑃𝑃 of a cell falling into the ‘Always’ category if each of the 9 iteration of ***S*** is assumed to be a fully independent event can be determined as 𝑃𝑃 = (0.2731)^9^ and would be expected to occur in 1 out of every 188,333 cells (0.000845%). The probability of falling into the ‘Never’ category assuming that recruitment at each iteration is independent is 𝑃𝑃 = (1 − 0.2731)^9^ or 1 out of ∼18 cells (5.66%). In contrast, out of 2,189 cells observed across the 9 iterations of ensemble ***S*,** Always cells represented 0.18% of the population, a 213 times greater proportion than would be expected if ensemble recruitment was unweighted across the population. Never cells represented 8.82% of the population, 1.6 times more than expected without weighting. Though the total number of cells categorized as Always and Never were very few, just 4 and 124 respectively, the observed distributions differed markedly from shuffled controls **(Figure S3A-E).** These results suggest that although the large majority of ensemble members are stochastically recruited to participate in a given iteration, there is a small minority of core members which contribute to all iterations of the ensemble.

For the ensembles evoked by stimulations with increasing train length, where the mean recruitment probability over the 5 iterations was 0.4393 (**Figure 3I**), Always cells represented 6.01% of the population, 3.7 times greater than expected if no weights were applied, while Never cells represented 11.11% of the population, 2 times greater than expected without weighting. Statistical comparisons of the number of cells in each category with each of the 10,000 reshuffled controls revealed that the observed categorizations were indeed consistently different than would be expected by chance **(Figure S6B-D).** To determine whether weighting of recruitment probability was related to cells’ contribution to the ensemble, we performed the same analysis when only excited cells were considered members of the ensemble (termed excited subassembly) or when only inhibited cells were considered members of the ensemble (termed inhibited subassembly) (**Figure 3K**). We found that, across both datasets, within the excited and inhibited subassemblies the proportion of the population classified as Always was reduced or eliminated entirely, demonstrating that the majority of Always cells in the full ensemble were recruited through both excitatory and inhibitory activity over iterations (**Figure 3L**-**O**). In contrast, the proportion of Never cells was greatly increased in both subassemblies, such that the overall distributions of Always, Sometimes, and Never cells within the subassemblies were consistently different than their respective shuffle controls **(Figure S3F-M)**. Together, these results further support that although the large majority of ensemble members are labile, there are core members that are consistently recruited and participate in all iterations of a given ensemble. Even within the core members however, the contribution to the ensemble remains labile with cells displaying excited and inhibited responses across iterations. Further, it appears that while the microcircuitry is organized such that most cells are recruitable to any given network, a minority are excluded or have a very low probability of being recruited by a given input.

Finally, we sought to determine whether individual differences in behavioral metrics of cognitive performance were related to microcircuit excitability observed in isolated dlPFC *ex vivo* and, if so, whether these relationships would be reflected in readouts of ensemble membership size or in the dynamics of population level signals. All subjects from the recordings presented thus far had gone through extensive behavioral testing which included attentional set-shifting (**Figure 4A**). Based on prior characterization of this task ^31^, three measures of individual performance were selected *a priori* for analyses: the median latency to touch the screen over sessions, change in performance over sessions as measured by the slope of the performance index (P_I_) across sessions, and the mean perseverative error rate across sessions (**Figure 4B**-**D**). To probe for latent relationships between single-cell dynamics and cognitive flexibility, we correlated these three behavioral metrics of interest with the proportion of the population excited across increasing stimulation intensities. Across all three behavioral metrics, we found no correlation with the proportion of the population excited by any of the stimulation intensities tested (**Figure 4E**-**H**, **Figure S4A-C)**. Next, we examined the relationship between the amplitude of the population-level response, measured as the peak amplitude of the whole field bulk fluorescence, and these same behavioral parameters (**Figure 4I**). No relationships were detected between median latency to touch and the peak amplitude (**Figure 4J**). Interestingly, we found the slope of the P_I_ was positively correlated with the peak evoked amplitude by stimulations with intensities of 4 mA and higher, demonstrating that greater microcircuit excitability is associated with higher cognitive performance (**Figure 4I**-**L**, **Figure S4D-F).** Further, the average perseverative error rate was negatively correlated with peak amplitudes evoked by high stimulation intensities (**Figure 4I**-**L**). Thus, our data directly link the intrinsic features of dlPFC microcircuits, observable in an isolated preparation, with higher-order cognitive functions and suggest that the ability to produce similar population level signals from predominantly distributed, labile networks is a key feature of dlPFC function.

**Figure 4.**
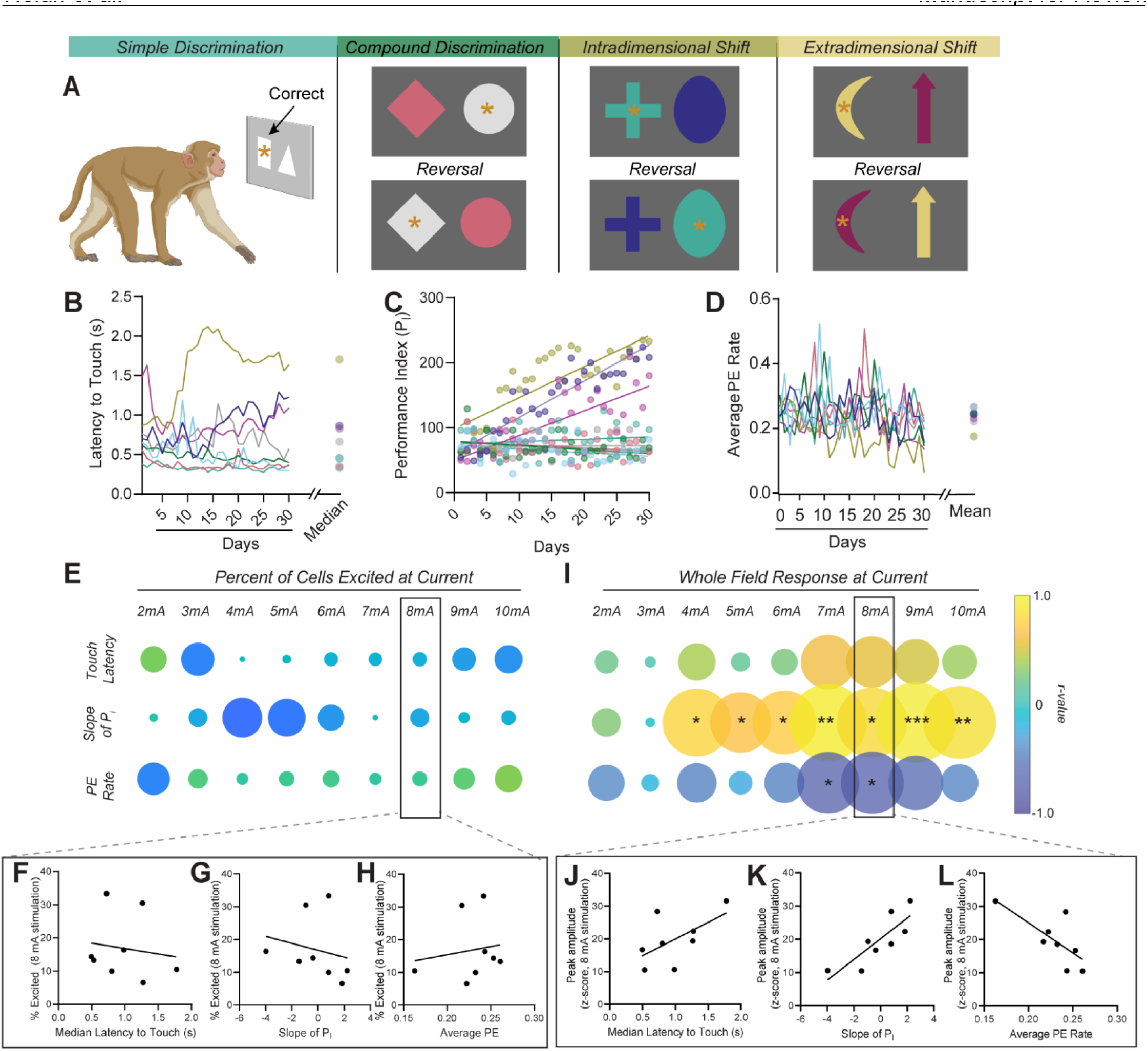
Cognitive flexibility is associated with intrinsic excitability of dlPFC microcircuits. **(A)** Diagram of set-shifting behavioral paradigm. Monkeys were trained to select one of two geometrical shapes, which would result in a food pellet reward, and as the animals reached the performance criterion they would advance to the next, more challenging phase (i.e., Simple Discrimination → Complex Discrimination). Each session was 45 minutes, and began with Simple Discrimination, regardless of the previous session’s performance. Three behavioral metrics of interest were selected *a priori* for analyses based on prior characterization of the task: median latency to touch the screen, slope of the performance index (P_I_), and mean perseverative error rate. **(B)** Latency to touch the screen from trial onset for each of the 8 subjects over 30 sessions. While there is session to session variance a clear pattern of individual differences can be seen, which are well described by the median latency over sessions. **(C)** Overall performance, as measured by the performance index (P_I_) over sessions. There are wide individual differences in performance over days, which can be summarized by the slope of the linear regression. Subsequent correlations between neural data and the rate of learning were performed using slope as a metric of learning rate. **(D)** Average perseverative error rate per session. A perseverative error refers to continued utilization of the previous strategy in a new phase of the task. Perseverative error rate (defined as perseverative errors as a proportion of the total number of errors) from each session was averaged together per subject and this value was used for correlation analysis. **(E)** Pearson’s correlation coefficients resulting from correlation analysis between the three behavioral metrics and the percent of the population excited by stimulations of increasing intensity (r values are indicted on the color axis and absolute r value is indicated by bubble size). No correlations were found across any of the variables tested. **(F-H)** Panels F, G, and H depict the scatter plots of data for the 8 mA stimulation with each of the three behavioral metrics (median latency: Pearson’s r = -0.15, p > 0.05; slope of P_I_: Pearson’s r = -0.21, p > 0.05; average PE rate: Pearson’s r = 0.16, p > 0.05). **(I)** The amplitude of the whole field response evoked by high stimulation intensities is positively correlated with the slope of the P_I_ and negatively correlated with average perseverative error rate. **(J-L)** Scatter plots and linear regressions depicting the relationships between the amplitude of the whole field response evoked by the 8 mA stimulation and each of the three behavioral metrics. **(J)** Median latency to touch (Pearson’s r = 0.59, p > 0.05). **(K)** Slope of the P_I_ (Pearson’s r = 0.83, p < 0.05). **(L)** Perseverative error rate (Pearson’s r = -0.73, p < 0.05). **\*,** p < 0.05, **\*\*,** p < 0.01

In summary, we have established a viable approach for the observation of large-scale population dynamics in *ex vivo* non-human primate brain tissue, circumventing long-standing technical limitations, and thereby revealing inherent properties of the dlPFC. We demonstrated that the intrinsic architecture of the dlPFC is capable of propagating signals through recurrent connections that can be effectively shaped into distinct representations of a variety of stimulus features. Our findings further indicated that activity in response to discrete inputs propagates through largely disparate and flexible networks of cells, supporting the wealth of evidence for distributed population representations across a cortical region, but suggest that a small minority of core members in the network may play specialized or essential roles. Finally, we revealed covariation between the static features of intracortical excitability observed *ex vivo* and individual differences in cognitive performance, thus linking cognitive abilities with microcircuit properties in an evolutionarily specialized region of PFC specific to primates. These results contribute to the understanding of intracortical microcircuitry and its relevance to cognitive processing, providing a foundation for further investigations into the synaptic mechanisms underlying cognitive performance.

**Supplemental Video S1. Representative field of view depicting spontaneous activity in the absence of extrinsic inputs and exogenous stimulation.** Video depicts a representative field of view, displayed in raw pixel intensities, with no stimulation delivered. Video playback is at 4x real time. Fluo-4 AM loaded cells are readily visible throughout the field of view and spontaneous fluorescence / calcium transients are apparent in many cells.

## Materials and Methods

### Subjects

Nine rhesus macaques (*Macaca mulatta*, five female and four male) averaging 6 years of age were used for the current study. Animals were individually housed in quadrant cages (0.8 x 0.8 x 0.9m) with constant temperature (20 – 22 degrees C) and humidity (65%) and an 11-hour-light cycle (lights on at 8 am). Animals were housed in the same room with visual and auditory access to all subjects in the room. In addition, monkeys were socially housed with a same-sex “partner” monkey for at least 2 hours per day or provided grooming contact if full pairing was contraindicated.

Each housing cage had a computer-operated panel embedded in the side wall that provided all food, fluids, and cognitive task presentations (additional details in ^31,32^). Body weights were taken weekly. All animal procedures and experiments were approved by the Oregon National Primate Research Center (ONPRC) Institutional Animal Care and Use Committee and followed recommendations of the *Guide for the Care and Use of Laboratory Animals* (National Research Council <u>2011</u>). Additionally, ONPRC is fully accredited by the Association for Assessment and Accreditation of Laboratory Animal Care International.

### Tissue extraction and slice preparation

Tissue was prepared as described previously ^18,19^. Briefly, subjects were deeply anesthetized with ketamine (10 mg/kg) and maintained on isoflurane. Monkeys were then perfused with ice-cold oxygenated perfusion solution [containing (in mM) 124 NaCl, 23 NaHCO_3_, 3 NaH_2_PO_4_, 5 KCl, 2 MgSO_4_, 10 D-glucose, 2 CaCl_2_]. Next, brains were removed quickly and sectioned along the coronal plane using a brain matrix (Electron Microscopy Sciences, Hartfield, PA, USA). The brain knife position was preplanned according to each subject’s prior MRI results. An isolated tissue block containing prefrontal area 46 was placed in ice-cold oxygenated perfusion solution on ice for subsequent transport between where monkeys were housed and the building where slicing was conducted.

A ceramic blade was attached to a vibrating tissue slicer to prepare 250 µm thick coronal brain sections containing area 46 of the dorsolateral prefrontal cortex as previously described ^19,33,34^. Tissue was sliced and rested for a minimum of 1 hour in oxygenated artificial CSF (aCSF) containing the following in mM: 126 NaCl, 2.5 KCl, 1.2 NaH_2_PO_4_, 2.4 CaCl_2_, 1.2 MgCl_2_, 25 NaHCO_3_, 11 D-glucose, and 0.4 L-ascorbic acid (pH adjusted to 7.4).

### Slice Incubation and Dye Loading

Once prepared, slices were divided into floating mesh-bottom wells which held the slices just below the surface of the aCSF reservoir which was held inside a lighttight incubation chamber (Braincubator, PAYO Scientific, Kingswood, AUS) ^35^. The incubation chamber consisted of a Peltier plate which held the reservoir, temperature and pH probes, aerator delivering 95%/5% carbogen, and a recirculating pump. Temperature was maintained at 15 degrees C and pH was adjusted back to 7.4 if any drift occurred. To maximize slice viability, aCSF was continuously removed from one side of the reservoir and returned on the other after passing through a UVC irradiation filter in a secondary chamber which was optically isolated from the slices. This ensures a constant supplied of oxygenated, clean aCSF and leads to extended slice viability ^36,37^. Slices were incubated in this chamber for at least 1 hour before dye loading.

After a minimum of 1 hour in the incubation chamber post-slicing, slices were removed and loaded with 5 µM fluo4 AM. Dye solution was made by dissolving lipophilized fluo-4 AM in pure DMSO containing 1% w/v pluronic acid (made fresh within the hour) to a concentration of 5 mM. Based on extensive parameterization of dye loading protocols from Yuste and colleagues ^38,39^, the dye solution was pipetted directly onto the surface of the slice such that a high concentration was briefly achieved before rapidly diffusing into the chamber and equilibrating to the final concentration. 4 µL of dye solution was pipetted on top of the slice which was held in in a low-volume oxygenated loading chamber containing aCSF (Brain Slice Keeper, Scientific Systems Designs Inc, Ontario, CAN). The total volume in the loading chamber was 4 mL, thus final concentrations were: 0.1% v/v DMSO, 0.001% w/v pluronic acid, and 5 µM fluo4 AM in aCSF. Slices were protected from light throughout the process. Following loading, slices were rinsed briefly in dye-free, pre-oxygenated aCSF before transferring them back to the incubation chamber. Slices were allowed to rest for at least 30 minutes before transferring to the imaging system.

### Overview of Imaging System

The system used for recording *ex vivo* calcium dynamics consisted of a customized upright laser scanning confocal microscope (Thorlabs Imaging Systems, Sterling, VA) integrated with a slice perfusion chamber (Warner Instruments, Hamden, CT). While in the imaging chamber, the flow rate was monitored online using pass-through flow sensors (Sensirion Inc, Chicago, IL) to ensure a stable supply of dissolved oxygen was delivered to the slice and that pharmacological agents were delivered in a reproducible manner. To ensure microsecond repeatability of synchronization between image capture and stimulation, the microscope and stimulus isolator (NL800A Current Stimulus Isolator, Digitimer, Hertfordshire, UK) were triggered using one function generator with a delay line such that both systems were controlled by a single master clock (PulsePal, Open Ephys, Rochester, NY). Fluorescence excitation was achieved with a 488 nm laser line fiber-coupled to the scan head. Emission was collected through a pinhole confocal with the imaging plane, and passed to a PMT detection module through a multimode fiber. Emission was then split by a 550nm longpass dichroic mirror and the reflected beam passed through a 525nm emission filter at the face of a cooled GaAsP PMT sampled at 80 MHz.

### Image Acquisition

Following a minimum 30-minutes rest period after dye loading, slices were transferred to the imaging chamber and immersed in aCSF which was flowing at a rate of 1 mL/min. For evoked recordings, a 4x magnification objective (4X Olympus Plan Achromat Objective, 0.10 NA, 18.5 mm WD, Thor Labs, Newton, NJ) was first utilized to assess gross anatomical features and place the stimulating electrode before switching to a 16x water dipping objective (16X Nikon CFI LWD Plan Fluorite Objective, 0.80 NA, 3.0 mm WD, Thorlabs, Newton, NJ) for the duration of the recordings.

During experimental recordings, images were acquired at 1024 x 1024 pixel density across a 762µm^2^ field of view (744 nm^2^ pixel size) at 15.14 frames per second (fps). For evoked recordings, stimulation onset was 5 seconds from the start of the frame capture and activity was recorded for 14.82 seconds following stimulation onset (19.82 seconds per recording, corresponding to 300 frames). The resonance scanner was kept constantly running during the inter-stimulation interval to reduce thermal drift.

### Electrical and Pharmacological Manipulations

Localized electrical stimulation was delivered via a twisted bipolar electrode (0.127mm diameter stainless steel 2-channel electrode, PlasticsOne, Roanoke, VA) held by a micromanipulator (MM-3 Micromanipulator, Narishige, Amityville, NY) and positioned superficially with contacts parallel to the principal sulcus. Calcium dynamics were evoked using a variety of stimulation parameters delivered to the slice. To determine the input-output relationship with varying stimulation amperage, we delivered a single pulse ranging from 2-10 mA, 4 ms wide, monophasic, constant current. The stimulation intensities selected here are within the range used in clinical application of intracortical stimulations ^40–42^. For the multi-pulse data, pulse intensities were held constant at 5 mA, and consisted of 4 ms wide monophasic pulses with train lengths increasing from 1 to 20 pulses. For the calcium chelation experiment, aCSF containing 10 µm BAPTA-AM (CAS: 126150-97-8, Tocris Biosciences, Minneapolis, MN, USA) was perfused onto the slice at a rate of 2 mL per minute for at least 30 minutes after the baseline recordings were taken.

### Calcium Imaging Analysis

#### Image processing and bulk fluorescence signal extraction

Following the acquisition, tiff frames were collated into one file for each sweep/recording using customized FIJI macros. Videos were then visually inspected and motion corrected, if needed, in Inscopix Data Processing software (Inscopix, Mountain View, CA). The bulk fluorescence signal, or whole field activity, was determined by averaging pixel intensities over the entire field of view to create a single brightness over time trace for each video, expressed in units of ΔF/F calculated as (F-F_0_)/F_0_ for each sample in the trace F and F_0_ equal to the mean intensity of the entire trace. To quantify the stimulation evoked activity, traces were z-score normalized to the first 70 frames (the pre-stimulation period spanned from frame 1:75), and the traces were then transformed such that the value of the last frame before the stimulation was set to zero and this differential was applied to the entire trace. Parameters of interest, peak amplitude following stimulation and area under the curve of the decay phase were then calculated using custom MATLAB scripts.

#### Single cell demixing and quantification

After the steps listed above, videos that were processed for whole-field analysis were then co-registered across sweeps using PatchWarp ^43^ to ensure that cells could be tracked effectively across the time series. Following co-registration, videos were then frame averaged by a factor of 2 bringing the final framerate to 7.58 fps and binned by a factor of 2 bringing the final pixel size to 1.49 µm^2^. We then used a constrained non-negative matrix factorization (CNMF) algorithm to extract fluorescence traces from cells within the field of view. All spatial footprints were then visually inspected following CNMF and any overlapping cells or cells that contained multiple cells in one footprint were removed.

For stimulus-evoked activity, representative heatmaps were created using traces that had been z-scored to a pre-stimulus window (−32 through -7 frames prior to the stimulation onset, roughly corresponding to 1 after the start of the recording through 1 second prior to the stimulation). Cells were classified as excited, inhibited, or no response based on the area of the curve during the 2 seconds following the stimulation onset. On a given sweep, cells with an AUC of above 0.85 were classified as excited while cells with an AUC of below 0.85 were classified as inhibited. These thresholds were empirically verified as effectively separating subpopulations and control experiments confirmed that both excited and inhibited responses reflected stimulation-evoked calcium dynamics (Figures S4 and S5).

### Attentional Set-Shifting Task (ASST)

#### Apparatus

For the set-shifting experiments, a computer-operated touchscreen panel was embedded in a side wall of each housing cage as previously described ^31,44^ and individual computers were controlled and inputs time-stamped using a centrally located computer. In addition to the touchscreen, there was a dowel below the screen, two drinking spouts on the screen’s left and right sides, and an infrared finger-poke on the righthand side (Med Associates Inc, US). These operanda were connected to a food pellet dispenser that delivered a 1g banana-flavored food pellet as a reward. All panels were linked to a network and the main computer via a National Instruments interface and Labview software (LabView 2011, SP1, National Instruments, Texas, United States).

#### Task design

As described in Figure 3, all animals underwent set-shift testing, which has been examined extensively and described previously for other cohorts ^31,44^. Briefly, animals were given the set shift sessions inside their housing cage each morning, alongside all other monkeys in the cohort. To establish touching the screen, highly preferred photographs (e.g., fruit, other monkeys) were projected and, when touched, expanded to the full screen for 3 sec ^31^. Following the acquisition of a reliable operant response (i.e., touching the screen), set-shifting training began. During training, the monkeys were presented with a random presentation of two geometrical shapes, and the trial ended following the selection of one of the shapes or after 30 seconds elapsed. Discrimination between which shape was predetermined as “correct” or “incorrect” was achieved by the contingent presentation of a 1g banana flavored food pellet and a presentation of the preferred photograph for a correct response and the contingent presentation of a blank screen for 10 seconds following an incorrect response. Subjects were gradually moved to a second order of FR1 (FR3) where every correct response earned a presentation of the preferred photo and every 3^rd^ correct response resulted in a pellet, to prevent satiety over the session. The criterion for the training phase was that responding was maintained throughout the 45-minute session. All animals received 7 training days on the second-order schedule.

Following training, animals were given 30 consecutive sessions to assess set-shifting performance followed by an additional 10 days at a later point. All behavioral variables were calculated by taking an average of the performance at the two timepoints. Within each session, there was a maximum of 4 original types of discrimination sets – simple, compound, and intradimensional and extradimensional shifts (shown in Figure 3A) each followed by a reversal set. A set consisted of a pair of shapes and each set retained the same two shapes and colors for all trials. The color of each shape and the side (right or left of center screen) of the screen it was presented was randomly chosen in each trial. Reversal trials included the same two shapes and colors with only the correct contingency being reversed. A set was considered acquired when a running twelve out of 15 consecutive trials was deemed correct. In simple discrimination, both shapes were either black or white, therefore only shape could be used as the basis of the discrimination. In the compound discriminations, the shapes were new as were the colors of the shapes, with the discrimination still based on the shape of the object. In the intradimensional shift set, there was a different combination of shapes and colors, and the discrimination was based on shape again. The fourth and final set represented an extradimensional shift, presenting a different combination of unique colors and shapes but with the correct discrimination now based on color. Progression through the sets was self-paced and the overall session ended after 45 minutes elapsed or criteria being met for all eight sets in the allotted time.

#### Behavioral data analysis

Based on previous work, the main set-shifting variables were as follows: the performance index (see ^31^ for details), the median latency to touch the screen, and the average ratio of the perseverative errors to total errors (non-reinforced stimuli selection). Perseverative errors were defined as errors during the reversal phase, where choices were based on the prior contingency ^45^ All variables were examined as a composite measure across the first 30 days and the second 10-day re-test.

### Statistical Analyses

All details regarding individual statistical comparisons can be found within the relevant figure legends. Statistical comparisons were conducted using Graphpad Prism 9 Software (Boston, MA, USA) with the exception of the permutation/reshuffling analyses in Figure 2 and Figure S6, which was performed using custom MATLAB scripts.

## Acknowledgments

This work was supported by NIH grants R00 DA04510 (NIDA), R01 AA030115 (NIAAA), U01 AA029971 (NIAAA), Alkermes Pathways Research Award, the Brain Research Foundation, the Whitehall Foundation (C.A.S), and P51 OD011092 (ORIP, K.A.G.). S.O.N. was supported by an NIH fellowship (F32DA051136). P.R.M. was supposed by an NIH fellowship (F31AA029626).

