## Supplemental Figures for "Recurrent activity within microcircuits of macaque dorsolateral prefrontal cortex tracks cognitive flexibility"

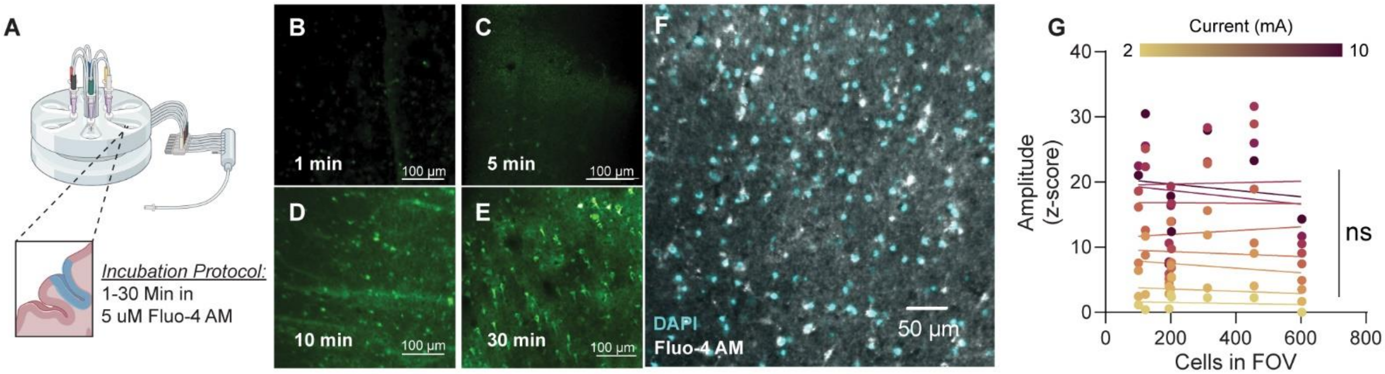


**Supplemental Figure S1. Time course of fluo-4 cell loading in acute slices of rhesus macaque area 46.** To conserve the number of animals used, this optimization was performed qualitatively and therefore does not include any statistical claims. **(A)** To optimize the loading protocol, we varied the loading period from 1 to 30 minutes on separate slices followed by a 30-minute rest period before imaging. **(B)** At 1 minute loading period, there was a minimal amount of cell loading, limited to small puncta-like staining. **(C)** After 5 minutes, there is noticeably more neuropil staining as well as more defined cell bodies, though this remained limited. **(D)** After a 10-minute loading period, there was noticeably more cell-like staining as well as neuropil staining, but this was further improved after **(E)** a 30-minute loading period where cell processes can be visualized. **(F)** Colocalization of fluo-4 AM loaded cells with post-hoc DAPI staining. **(G)** Scatter plot and linear regressions of the relationship between the amplitude of whole field responses evoked by each of the stimulation intensities and the number of cells segmented in the field of view. There was no correlation found for any of the stimulation intensities tested: (all r values refer to Pearson's correlation coefficient; 2 mA: $r_{(8)} = -0.13$, $p > 0.05$; 3 mA: Pearson's $r_{(8)} = -0.27$, $p > 0.05$; 4 mA: $r_{(8)} = -0.18$, $p > 0.05$; 5 mA: $r_{(8)} = -0.08$, $p > 0.05$; 6 mA: $r_{(8)} = 0.08$, $p > 0.05$; 7 mA: $r_{(8)} = -0.008$, $p > 0.05$; 8 mA: $r_{(8)} = 0.03$, $p > 0.05$; 9 mA: $r_{(8)} = -0.14$, $p > 0.05$; 10 mA: $r_{(8)} = -0.11$, $p > 0.05$;). FOV = field of view; ns = not significant, $p > 0.05$.

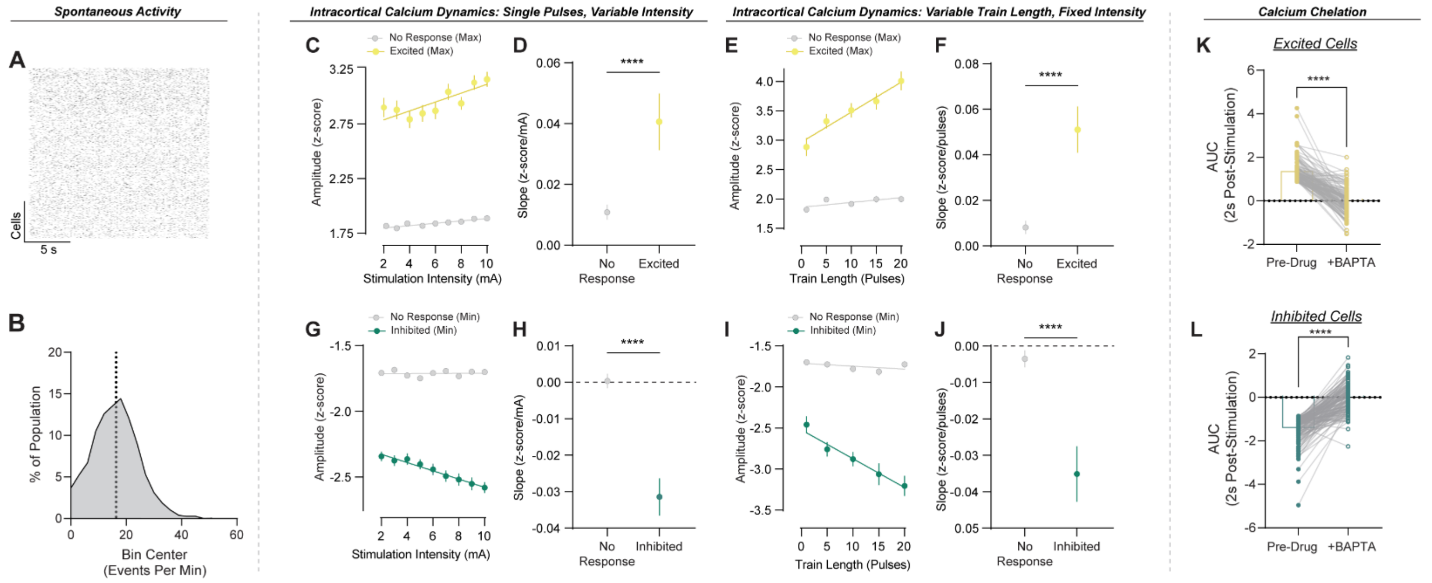


**Supplemental Figure S2. dlPFC microcircuitry intrinsically supports a high rate of spontaneous activity and exhibits diverse responsivity to uniform inputs.** Event detection was used to identify transients over time in the absence of stimulation. **(A)** Raster plot of events detected across individual cells from a 20 second recording with no stimulation. **(B)** Frequency distribution of event rates per cell for 2,189 cells. The population mean across all cells was 16.37 events/minute (SEM: +/- 0.16) as indicated by the vertical line on the histogram, with a range from 0 to 51.91. **(C)** Linear fits of the maximum amplitude (in z-score) for cells classified as excited or no response. Both excited and no response cells' amplitudes increase over stimulation intensities (mA) (No Response slope vs 0: $F_{(1,14318)} = 21.91$, $p < 0.0001$; Excited slope vs 0: $F_{(1,2869)} = 19.6$, $p < 0.0001$). Though there is a relationship between amplitude and stimulation intensity in both cell groups, **(D)** the rate at which peak amplitude (z-score) increases over stimulation intensity (mA) is dramatically greater in excited cells compared to no response cells as measured by the slope of the linear regression (z-score/mA) (unpaired t-test$_{(1,17189)} = 4.52$, $p < 0.0001$). **(E)** Similar to the relationship with stimulation intensity, both excited and no response cells show a increase in peak amplitude (z-score) as train length increases (slopes differ from zero - Non-Responsive: $F_{(1,1537)} = 8.96$, $p < 0.01$; Excited: $F_{(1,768)} = 26.4$, $p < 0.0001$). **(F)** Excited and no response cells show different rates of increased response (amplitude) over train length (pulses), as the slope (z-score/pulses) is greater in excited cells (t-test$_{(1,2307)} = 5.36$, $p < 0.0001$). **(G)** The inhibitory amplitude (z-score below 0) for cells classified as inhibited or no response were also fit with linear regressions. Inhibited cells show larger magnitude inhibitory responses as stimulation intensity increases (Inhibited slope vs 0: $F_{(1,2507)} = 40.08$, $p < 0.0001$) while no response cells do not (No Response slope vs 0: $F_{(1,14318)} = 0.84$, $p > 0.05$). **(H)** The slope (z-score/mA) was greater for cells classified as inhibited compared to no response (unpaired t-test$_{(1,16827)}$ = 6.56, $p < 0.0001$). **(I)** This effect is also seen over increasing train length, where inhibited cells show larger magnitude inhibitory responses as stimulation intensity increases while cells classified as no response did not (No Response: $F_{(1,1537)} = 2.604$, $p > 0.05$; Inhibited: $F_{(1,434)} = 21.84$, $p < 0.0001$). **(J)** Similarly, the slope (z-score/mA) was greater for cells classified as inhibited compared to no response in the train length dataset (unpaired t-test$_{(1,1973)} = 5.50$, $p < 0.0001$). **(K)** As a negative control, in separate slices 10 µM BAPTA-AM was washed onto the slice after establishing a pre-drug baseline. Heatmaps of all cells aligned to the stimulation (1 pulse, 5 mA, 4 ms pulse width at time = 0s) before and after washing on BAPTA-AM. Calcium chelation dramatically reduced the evoked response in excited cells (paired t-test$_{(1,124)} = 19.22$, $p < 0.0001$), and **(L)** attenuated the inhibitory response in inhibited cells (paired t-test$_{(1,146)} = 20.17$, $p < 0.0001$). Error bars indicate SEM. ****, $p < 0.0001$.

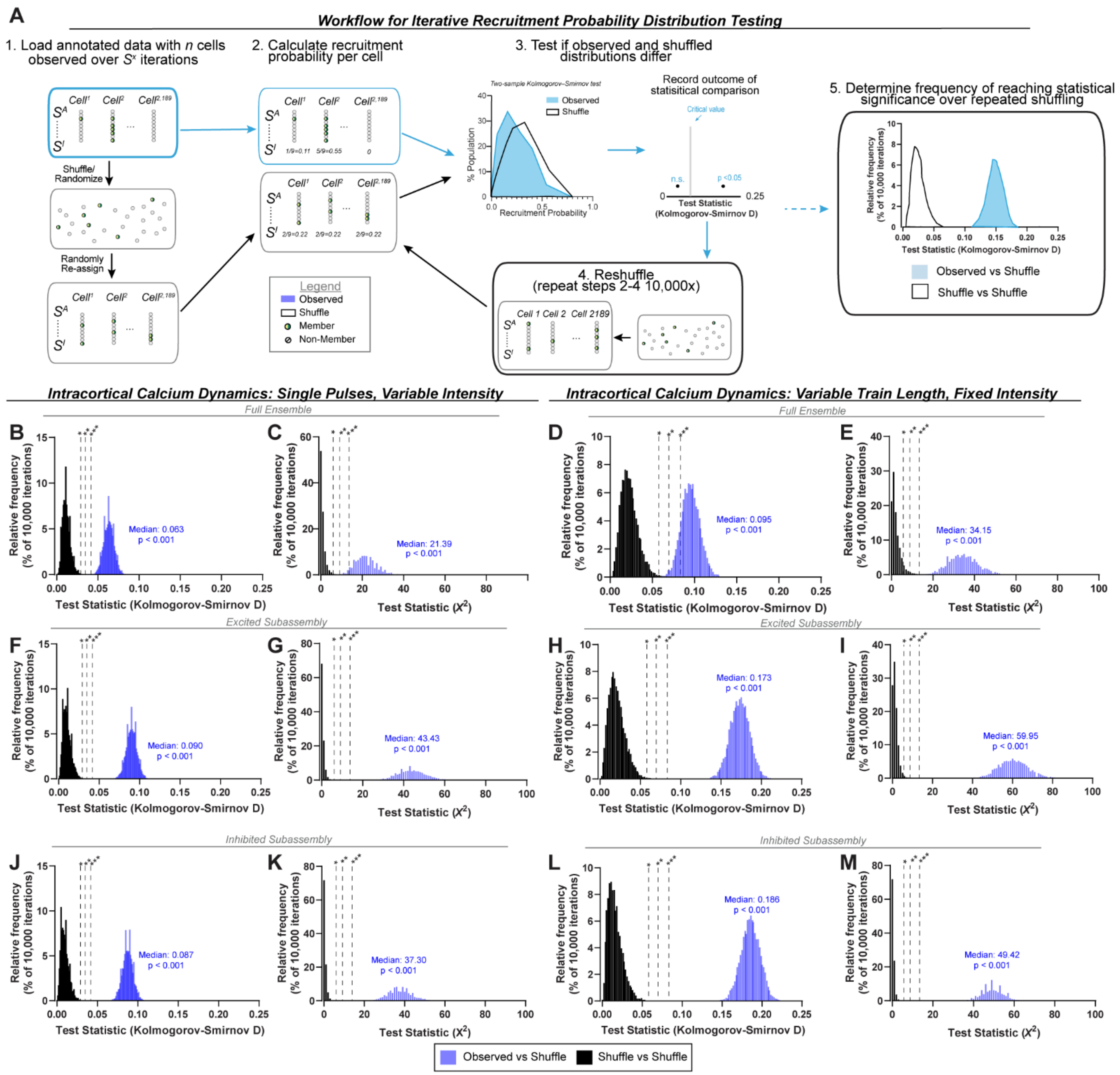


**Supplemental Figure S3. Permutation testing of ensemble recruitment probabilities.** Statistical comparisons supporting the claims made in Figure 3. **(A)** Diagram depicting the permutation pipeline used to compare observed patterns of stimulation-evoked responses to shuffled data. For clarity, the diagram depicts one of two datasets tested (the intensity curve dataset with 2,189 cells and 9 stimulations), and one of the two tests used (Kolmogorov-Smirnov test). The structure was identical for the pulse number dataset (549 cells and 5 stimulations) and for chi-squared tests (see below). **1.** A matrix was formed with columns equal to the number of cells in the dataset and rows equal to the number of stimulations. Each value in the matrix indicated whether the cell was part of the ensemble evoked by each of the stimulations. A shuffled dataset was then created by randomizing the location of values in the matrix. The observed and shuffled datasets thus had the same number of cells and stimulations, as well as the same total number of ensemble members, but any influence of cell identity on probability of ensemble membership over the stimulations was due to chance in the shuffled set. **2.** Recruitment probability was calculated for each cell in the dataset (number of times recruited / number of stimulations). **3.** Distributions of recruitment probability for the observed and shuffled datasets were compared using a two-sample Kolmogorov-Smirnov test, and the resulting test statistic (Kolmogorov-Smirnov D) was recorded. **4.** The shuffled dataset was reshuffled, and steps 2-4 were then repeated for 10,000 iterations. **5.** A distribution of the resulting 10,000 test statistic values was plotted, and the median value was used to determine significance. As a control to ensure statistical tests selected were appropriate and did not result in spurious error rates, this pipeline was repeated with both inputs as shuffled datasets. In the shuffle vs shuffle control, any significant outcomes detected are due to chance. For chi-squared tests, the pipeline was identical except for step 3 where instead of recruitment probability, each neuron in the observed and shuffled datasets was classified into one of three categories: Always (recruited by every stimulation in the dataset), Sometimes (recruited by at least one stimulation and not responsive to at least one stimulation), and Never (not recruited by any of the stimulations). The number of observations in each category was then compared between the observed and shuffled datasets using a chi-squared test for independence. The test statistic, $X^2$ in this case, was recorded and steps 2-4 were repeated as described above. **(B-M)** For each of the distributions, the critical values associated with alpha values of 0.05, 0.01, and 0.001 are indicated by vertical lines labeled as *, **, and ***, respectively. For Kolmogorov-Smirnov tests the critical values were determined as $D_{\text{critical}} = \frac{c(\alpha)}{\sqrt{n}}$ where $c(\alpha)$ is the critical value for alpha value $\alpha$ from the Kolmogorov distribution and $n$ is the sample size. For chi-squared tests of independence the critical values were determined from a chi-squared lookup table. All chi-squared tests performed had 2 degrees of freedom (number of categories [always, sometimes, never] minus 1), and therefore have the same critical values for all tests. **(B-E)** Results of the permutation tests when using the full ensemble as the input data (considering both excitatory and inhibitory responses as membership), corresponding to Figure 3A-E. **(B)** Distributions of Kolmogorov-Smirnov D values for the stimulation intensity curve when using the full ensemble (corresponds to Figure 3H). **(C)** Distributions of $X^2$ values for the stimulation intensity curve when using the full ensemble categorized into ‘always’, ‘sometimes’, and ‘never’ (corresponds to Figure 3H inset). **(D)** Distributions of Kolmogorov-Smirnov D values for the train length curve when using the full ensemble (corresponds to Figure 3J). **(E)** Distributions of $X^2$ values for the train length curve when using the full ensemble categorized into ‘always’, ‘sometimes’, and ‘never’ (corresponds to Figure 3J inset). **(F-I)** Results of the permutation tests when using the excited subassembly as the input data (considering only excitatory responses to the stimulation), corresponding to Figure 3L-N., diagram in Figure 3K. **(F)** Distributions of Kolmogorov-Smirnov D values for the excited subassembly in the stimulation intensity curve experiment (corresponds to Figure 3L). **(G)** Distributions of $X^2$ values for the excited subassembly categorized into ‘always’, ‘sometimes’, and ‘never’ in the stimulation intensity curve experiment (corresponds to Figure 3L inset). **(H)** Distributions of Kolmogorov-Smirnov D values for the excited subassembly in the train length curve experiment (corresponds to Figure 3N). **(I)** Distributions of $X^2$ values for the excited subassembly categorized into ‘always’, ‘sometimes’, and ‘never’ in the train length curve experiment (corresponds to Figure 3N inset). **(J-M)** Results of the permutation tests when using the inhibited subassembly as the input data (considering only excitatory responses to the stimulation), corresponding to Figure 3M and 3O, diagram in Figure 3K. **(J)** Distributions of Kolmogorov-Smirnov D values for the inhibited subassembly in the stimulation intensity curve experiment (corresponds to Figure 3M). **(G)** Distributions of $X^2$ values for the excited subassembly categorized into ‘always’, ‘sometimes’, and ‘never’ in the stimulation intensity curve experiment (corresponds to Figure 3M inset). **(H)** Distributions of Kolmogorov-Smirnov D values for the inhibited subassembly in the train length curve experiment (corresponds to Figure 3O). **(I)** Distributions of $X^2$ values for the inhibited subassembly categorized into ‘always’, ‘sometimes’, and ‘never’ in the train length curve experiment (corresponds to Figure 3O inset). ***,** $\alpha = 0.05$; ****,** $\alpha = 0.01$; *****,** $\alpha = 0.001$.

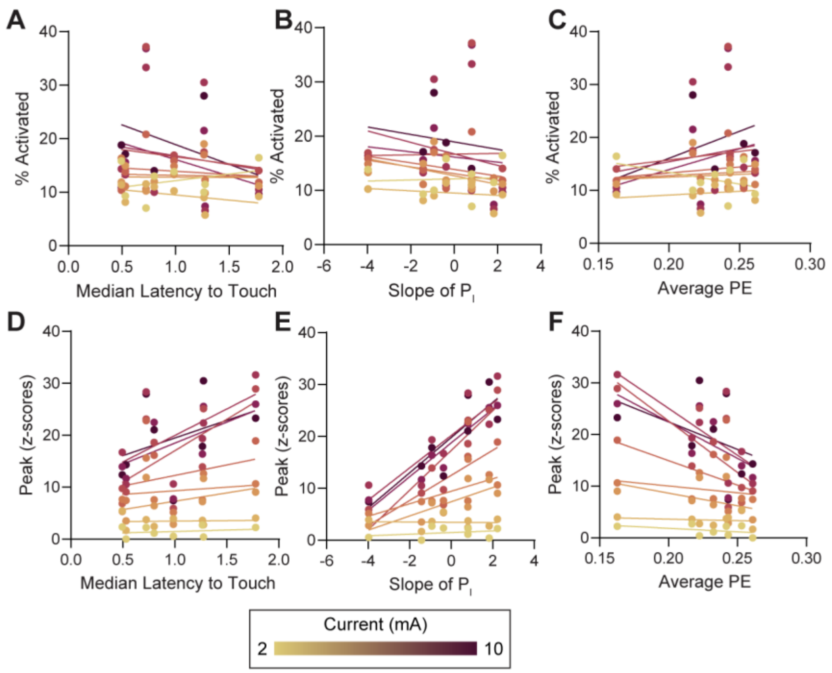


**Supplemental Figure S4. Bulk activity in far-field responses evoked by perisulcal stimulation covary with individual differences in set-shifting behavior**. **(A-C)** Scatterplots and linear regressions comparing set-shifting outcome measures [median latency to touch, slope of the performance index ($P_I$), or average perseverative error (PE)] to the proportion of cells activated by stimulations with 2 through 10 mA intensities, corresponding to the correlation coefficients reported in Figure 3E. **(D-F)** Scatterplots and linear regressions comparing set-shifting outcome measures to the peak amplitude of the whole field response evoked by stimulations with 2 through 10 mA intensities, corresponding to the correlation coefficients reported in Figure 3I.
